# Nitric Oxide Regulates the *Ralstonia solanacearum* Type 3 Secretion System

**DOI:** 10.1101/2020.10.26.355339

**Authors:** Connor G. Hendrich, Alicia N. Truchon, Beth L. Dalsing, Caitilyn Allen

## Abstract

*Ralstonia solancearum* causes bacterial wilt disease on diverse plant hosts. *R. solanacearum* cells enter a host from soil or infested water through the roots, then multiply and spread in the water-transporting xylem vessels. Despite the low nutrient content of xylem sap, *R. solanacearum* grows very well inside the host, using denitrification to respire in this hypoxic environment. *R. solanacearum* growth *in planta* also depends on the successful deployment of protein effectors into host cells via a Type III Secretion System (T3SS). The T3SS is absolutely required for *R. solanacearum* virulence, but it is metabolically costly and can trigger host defenses. Thus, the pathogen’s success depends on optimized regulation of the T3SS. We found that a byproduct of denitrification, the toxic free-radical nitric oxide (NO), positively regulates the *R. solanacearum* T3SS both *in vitro* and *in planta*. Using chemical treatments and *R. solanacearum* mutants with altered NO levels, we show that the expression of a key T3SS regulator is induced by NO in culture. Analyzing the transcriptome of *R. solanacearum* responding to varying levels of NO both in culture and *in planta* revealed that the T3SS and effectors were broadly upregulated with increasing levels of NO. This regulation was specific to the T3SS and was not shared by other stressors. Our results suggest that *R. solanacearum* experiences an NO-rich environment in the plant host and may use this NO as a signal to activate T3SS during infection.

## Introduction

*Ralstonia solancearum* (*Rs*) is a Gram-negative soil-borne betaproteobacterium that causes bacterial wilt disease on a wide range of plant hosts, including important crops like tomato, potato, and banana (Elphinstone 2005). The bacteria survive in and are spread through infected soil and water, infecting host roots through wounds and natural openings (Vasse et al. 1994). Once inside, *R. solanacearum* cells move to the xylem, where they can grow and spread throughout the plant. Although this bacterium is slow growing and sensitive to stress when grown *in vitro*, in plant xylem vessels *R. solanacearum* is a formidable force. *R. solanacearum* quickly grows to high cell densities in xylem even though this habitat is low in oxygen and nutrients, and accessible to host defenses (Genin and Denny 2012). As it grows, *R. solanacearum* produces an arsenal of virulence factors, including highly mucoid extracellular polymeric substances (EPS), a consortium of cell wall degrading enzymes, and dozens of protein effectors that are injected into host cells via the Type III Secretion System (T3SS) (Genin and Denny 2012). Water transport in infected plants is eventually blocked by a combination of host-produced gels and tyloses, degradation of vessel walls, EPS, and the sheer mass of bacterial cells. Without sufficient water, plants wilt and die while *R. solanacearum* cells escape back into the soil, where they can find another host (Swanson et al. 2007).

The lethality of bacterial wilt disease combined with its wide host range and a lack of effective treatments make *R. solanacearum* an important agricultural problem around the world (Yuliar et al. 2015; Elphinstone 2005). Successful disease management depends on understanding *R. solanacearum* biology, but the pathogen behaves very differently in culture than it does in its natural environment (Jacobs et al. 2012). In particular, the T3SS, which is absolutely essential for virulence and subject to a complex regulatory cascade, is affected by the plant environment in ways that remain poorly understood (Poueymiro and Genin 2009). The PhcA quorum sensing system upregulates T3 secretion in culture, but not *in planta* (Génin et al. 2005; Jacobs et al. 2012; Monteiro et al. 2012a; Khokhani et al. 2017). An unidentified ‘diffusible factor’ increases T3SS expression *in planta* via the HrpG global regulator (Genin and Denny 2012).

We have discovered that the diffusible free radical nitric oxide (NO) plays a role in *R. solanacearum* life inside plants. Xylem sap is rich in nitrate, which *R. solanacearum* uses as an alternate electron acceptor to respire in the hypoxic xylem environment (Dalsing et al. 2015). The *R. solanacearum* strain GMI1000 reduces nitrate completely to dinitrogen gas by means of denitrifying respiration. The genes required for this process are highly expressed during infection (Jacobs et al. 2012; Khokhani et al. 2017). However, denitrifying respiration produces NO as an intermediate. In large quantities, NO is toxic and strongly inhibits *R. solanacearum* growth. The pathogen manages this threat by detoxifying NO with the nitric oxide reductase NorB and the flavohemoglobin HmpX, which are both highly expressed *in planta* and required for full virulence (Dalsing et al. 2015; Jacobs et al. 2012). Plants use NO extensively for both signaling and defense, and can produce NO through multiple different pathways (Scheler et al. 2013; Delledonne et al. 1998; Simontacchi et al. 2015; Wilson et al. 2008; Leitner et al. 2009). Thus, *R. solanacearum* likely encounters NO during plant pathogenesis.

In this study, we identify a link between NO and regulation of the *R. solanacearum* T3SS. We first show that T3SS gene expression increases when *R. solanacearum* is actively denitrifying. We then show that this effect depends specifically on the presence of NO. Based on these results, we conducted a transcriptomics experiment to determine the effects of NO on *R. solanacearum* gene expression in culture and *in planta* and show that levels of many transcripts, including 107 T3SS-related genes, are altered by changes in NO levels.

## Results

### Nitric oxide changes expression of the *R. solanacearum* T3SS

Regulation of the T3SS in *R. solanacearum* is multi-layered and incompletely understood (Fig 1.A). Because *R. solanacearum* actively denitrifies *in planta*, we wondered if this pathway or its products affect T3SS expression during plant infection (Dalsing et al. 2015; Jacobs et al. 2012). To test this hypothesis, we used an *R. solanacearum* reporter strain carrying a fusion of the *lux* operon to the promoter of *hrpB*, which encodes a key T3SS regulator (Genin and Denny 2012). Transcription of *hrpB* increased >3-fold when the bacterium was denitrifying in culture (Fig. 1B).

**Fig 1.**
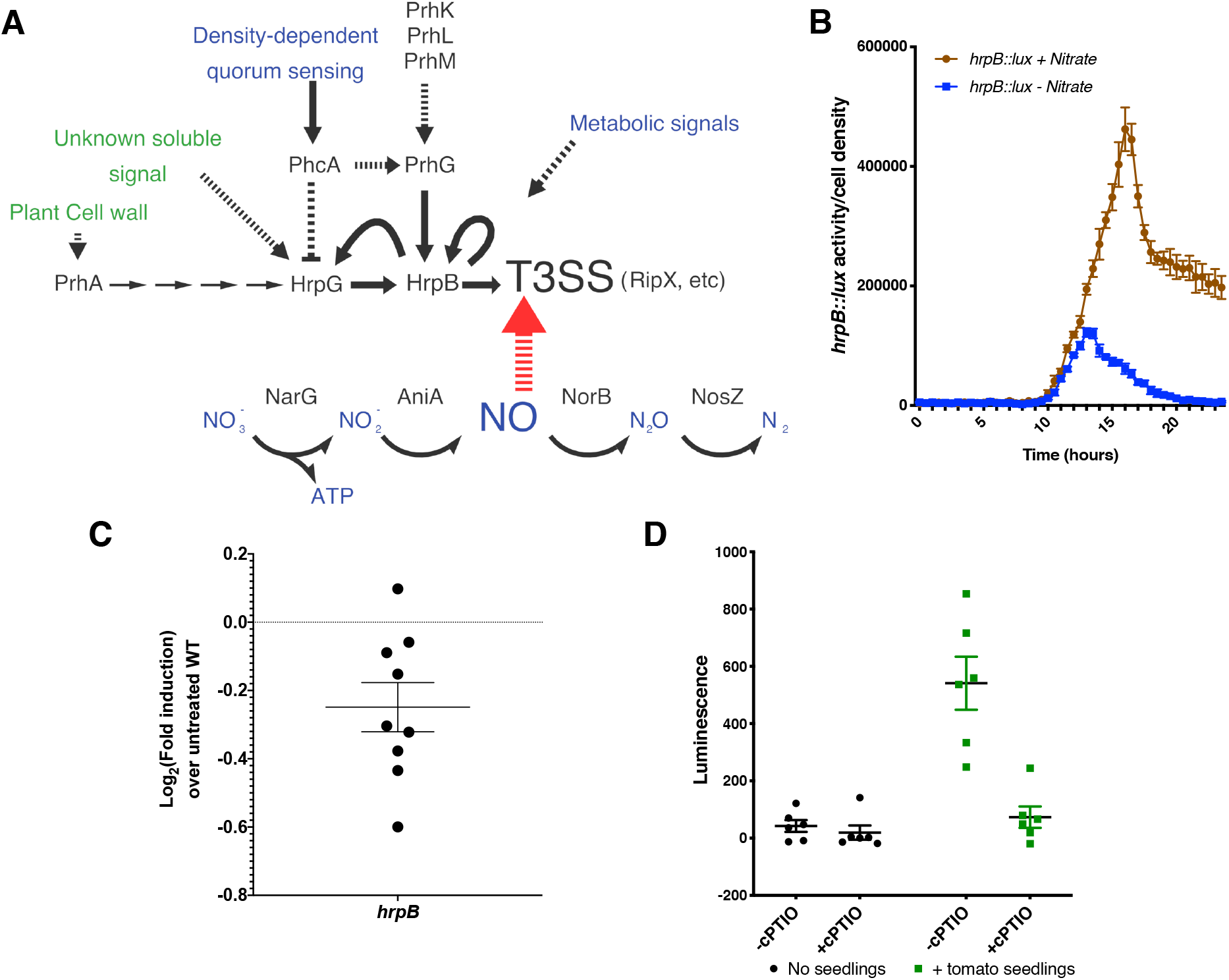
Nitric oxide affects expression of the *R. solanacearum* T3SS. **A**, Summary of known T3SS regulation in *R. solanacearum* strain GMI1000. Arrows indicate positive regulation, barred lines indicate repression, dashed lines indicate an unknown or uncharacterized mechanism, green text indicates plant-derived signals, and blue text indicates metabolites. T3SS genes, such as the T3 effector *ripX*, are controlled by the transcriptional regulator HrpB. HrpB is transcriptionally activated by HrpG and PrhG, which are themselves controlled by the quorum-sensing regulator PhcA, but only *in vitro*. PrhG activity also depends on three uncharacterized proteins, PrhK, PrhL, and PrhM, through an unknown mechanism. HrpG is activated by the contact-dependent plant cell wall sensor PrhA as well as by an unknown soluble signal present during plant infection. **B**, Expression of a bacterial luciferase transcriptional reporter of HrpB. *R. solanacearum* strain GMI1000 *hrpB::lux* was grown in denitrification-inducing conditions with or without 30 mM nitrate. Both culture density (OD_600_) and luminescence were measured every 30 min, and luminescence was normalized to culture density. Error bars represent standard error of the mean. Cultures grown with nitrate had significantly increased *hrpB* expression (2-way ANOVA, *P*=0.0003). **C**, The NO scavenger cPTIO reduces *hrpB* expression in *R. solanacearum* cells grown in denitrifying conditions. Cultures were grown as in B with and without 1 mM cPTIO and harvested at 16 hpi. *hrpB* expression was measured using qRT-PCR. Expression of *hrpB* was reduced in cPTIO-treated cultures (*P*<0.0001, students T test) Error bars represent standard error of the mean. **D**, The NO scavenger cPTIO decreased T3SS activity in the presence of a host. Luminescence of *R. solanacearum hrpB::lux* was measured after 24 h incubation in water with or without five day-old tomato seedlings, and with or without 1 mM cPTIO, an NO-scavenger. Exposure to tomato seedlings induced *hrpB::lux* expression (*P*=0.004, students T test). *R. solanacearum* cells exposed to both tomato seedlings and cPTIO had luminescence indistinguishable from that of cells without tomato seedlings. Error bars represent standard error of the mean.

An intermediate product of denitrification is the highly reactive free radical NO. Because NO is a common signaling molecule in eukaryotes, we decided to determine if it was responsible for *hrpB* activation observed during denitrification. To block the effects of NO, we used the NO-scavenging chemical cPTIO (Pfeiffer et al. 1997). While cPTIO is commonly used to both detect and deplete NO from solutions, it can also be reduced or inactivated by factors other than NO (D’Alessandro et al. 2013). To mitigate cPTIO effects unrelated to NO, we tested its ability to affect *hrpB* expression in two conditions. First, we incubated *R. solanacearum* in denitrification-promoting conditions with and without cPTIO. We measured *hrpB* expression in RNA extracted from these cultures at 16 hpi, near the peak of *hrpB::lux* expression. In these conditions, NO scavenging by cPTIO slightly but significantly reduced *hrpB* expression (Fig. 1C). We then tested the effect of a host on the interaction between *hrpB* expression and NO. We incubated a cell suspension of R. solanacearum *hrpB::lux* with tomato seedlings for 24 hours with or without cPTIO. As expected, the presence of a host plant activated *hrpB* gene expression. However, adding cPTIO to the mixture completely reversed this effect (Fig. 1D). Together, these results indicated that the denitrifying respiration-dependent increase in *hrpB* expression observed in culture is specifically triggered by NO. Further, this result suggests that NO could be the unknown soluble signal that increases *hrp* regulon expression *in planta*.

### Denitrifying metabolism causes broad changes in *R. solanacearum* gene expression

While the above results showed that NO can affect *hrpB* expression, many other genes are involved in regulation, biosynthesis, and function of the *R. solanacearum* T3SS. To better understand the broader effects of NO on pathogen biology and T3SS gene expression, we sequenced the transcriptomes of *R. solanacearum* cells grown with varying levels of NO both in culture and in infected tomato plants (Fig. 2A). In culture, we grew the cells in 0.1% O_2_ in a modified Van den Mooter medium (VDM), which promotes denitrifying respiration (Dalsing et al. 2015). We manipulated intracellular *R. solanacearum* NO levels genetically by using mutants lacking either the NarG nitrate reductase (the *ΔnarG* mutant cannot denitrify and does not produce NO), or the NorB nitric oxide reductase (the *ΔnorB* mutant accumulates NO, see Supplementary Fig. 1). Exogenous NO levels were manipulated by treating cells with CysNO, a nitrosothiol NO donor. Because high NO levels are toxic to *R. solanacearum*, we controlled for general stress responses by also profiling transcriptomes of cultures treated with H_2_O_2_, a non-NO source of stress. After about 18 h, growth of the *ΔnorB* mutant was inhibited by accumulating NO, so we sampled cultures 16 hpi, when *ΔnarG, ΔnorB*, and the wild-type strain were at similar densities (Supplementary Fig. 1). CysNO or H_2_O_2_ treatments were added 4 h before sampling, at the highest concentrations that did not significantly alter culture growth (Supplementary Fig 2A). Total RNA was extracted from bacteria *in planta* 3 days after tomato plants were inoculated with ~2000 CFU of *R. solanacearum* through a cut petiole (Khokhani et al. 2017). This method synchronized the infection process better than root inoculation. Bacterial colonization in sampled tissue was standardized by extracting RNA only from stems containing between 5×10^7^ and 10^9^ CFU/g stem, as determined by dilution plating a ground stem section (Supplementary Fig. 2B). Each of the three biological replicates contained pooled RNA from 4 or 5 plants.

**Fig 2.**
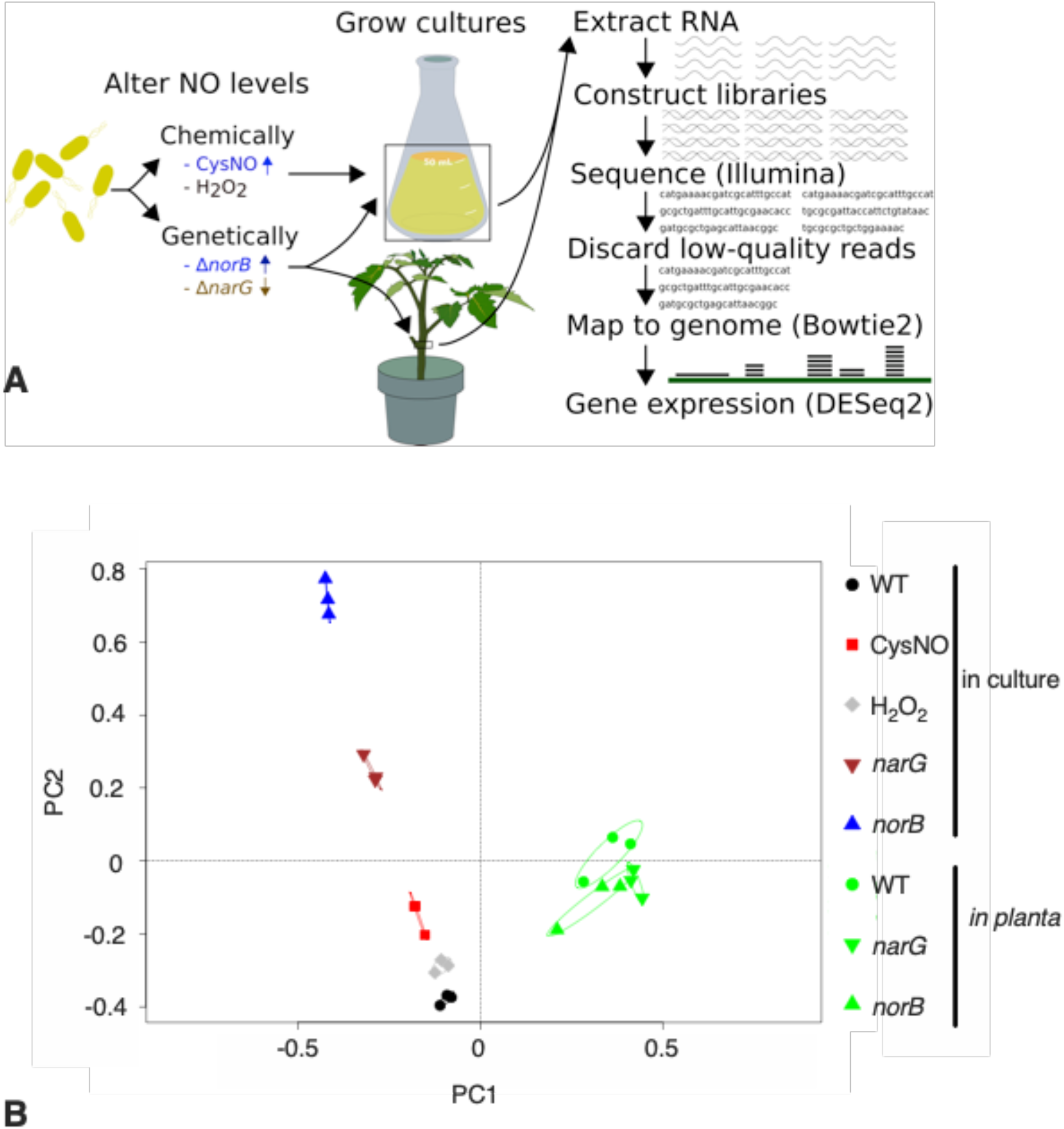
Nitric oxide causes broad changes in *R. solanacearum* gene expression. **A**, Design of RNA-seq experiment. We extracted RNA from *R. solanacearum* grown with altered NO levels from stems of infected tomato plants (*in planta)* and from denitrifying VDM medium (in culture) and NO levels were increased chemically by adding the NO donor CysNO to wild type (WT) and increased genetically by using a NO-accumulating *Δ*norB mutant. The *ΔnarG* mutant has reduced levels of NO. H_2_O_2_-treated WT was included as a control for non-nitrosative general stress responses. **B**, Whole-transcriptome datasets were used for principal component analysis; plot created in R with the package Vegan (Oksanen et al. 2019). Each data point represents one biological replicate, with three replicates per condition. Ellipses represent a 95% confidence interval for each condition (some ellipses are so small that they appear as lines).

Around 97% of RNA-seq reads from *in vitro* samples mapped to the *R. solanacearum* genome (Supplementary Fig. 3A). Because tomato stem slices also included host RNA, only 9.2% of the *in planta* RNA reads mapped to the *R. solanacearum* genome. However, the absolute number of bacterial transcripts from the *in planta* samples still exceeded 10^6^ reads per replicate. Principal component analysis clustered the *in vitro* samples together based on treatment, and the *in planta* samples were even more tightly clustered, reflecting less variation in gene expression than the culture samples (Fig. 2B). Genes were considered differentially expressed if they had an adjusted P-value <0.05. A full list of gene expression values can be found in the supplementary materials.

We validated our transcriptomic results by using qRT-PCR to measure the expression of two regulators of T3SS, *hrpB* and *hrpG*, and one well-expressed T3E, *ripX*. Expression levels of each of these genes in the *ΔnarG* and *ΔnorB* mutants generally matched those obtained from RNA-seq analysis (Fig. 3). As with *hrpB* expression, *hrpG* and *ripX* were both activated by NO, whether it was produced endogenously or exogenously.

**Fig 3.**
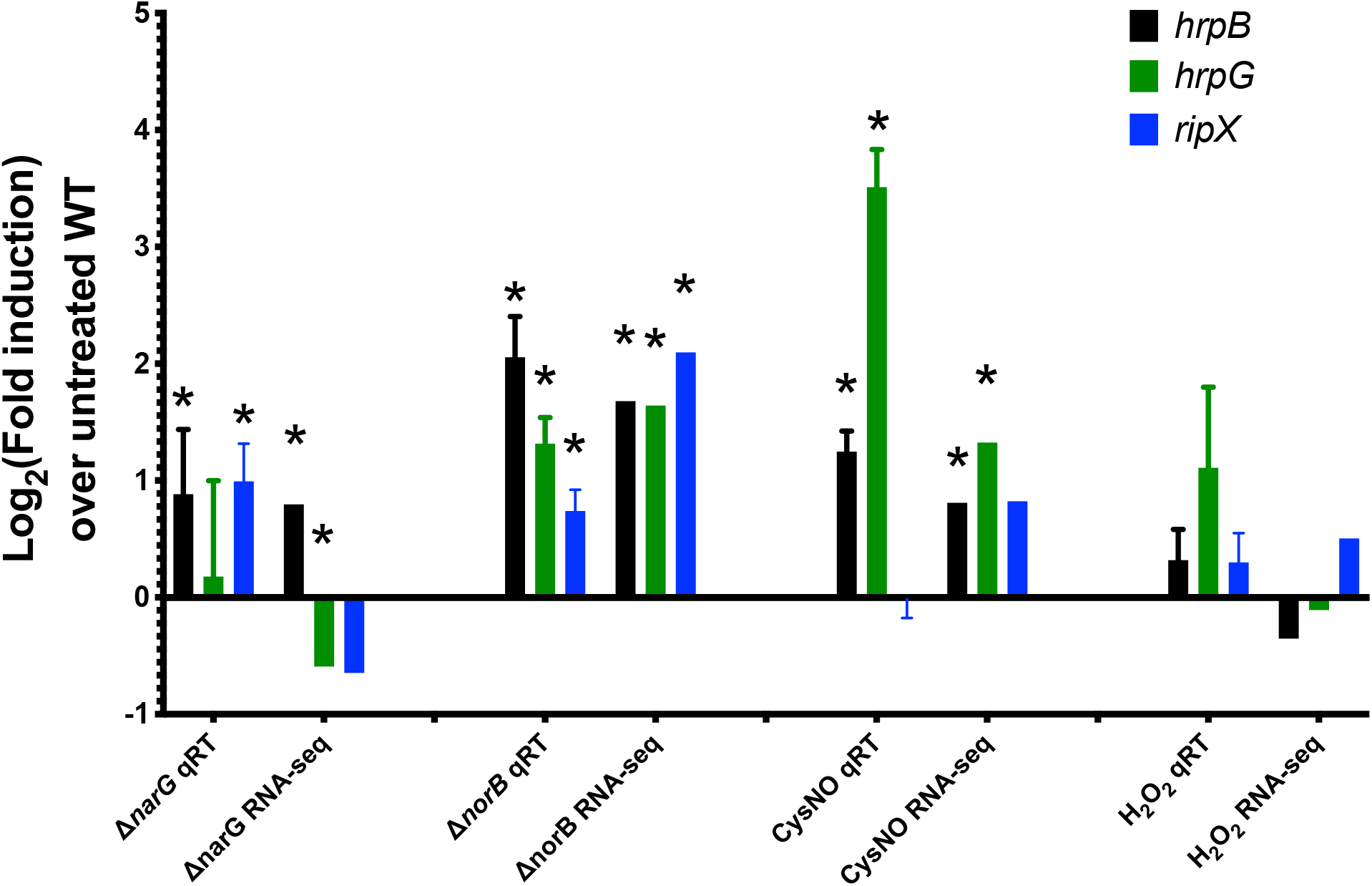
Expression analysis of selected T3SS-related genes by qRT-PCR validated RNA-seq results. RNA was extracted from *R. solanacearum* cultured in VDM media at 0.1% oxygen for 16 hours for both qRT-PCR and RNA-seq analysis. *R. solanacearum* samples were: wild type strain GMI1000 (WT, untreated control reference), non-denitrifying low-NO mutant *(ΔnarG*) and NO-accumulating mutant (*ΔnorB*), and WT treated for 4h with exogenous NO (CysNO). Gene expression levels in all samples were normalized using *serC* (Jacobs et al. 2012). Error bars indicate standard error of the mean. Black bars, log_2_fold-change expression of T3SS regulatory gene *hrpB*; green bars, log_2_ fold-change expression of T3SS regulatory gene *hrpG;* blue bars, log_2_ fold-change expression of Type 3-secreted effector gene *ripX*. * *P*<0.05, for qRT, student’s t-test compared to gene expression level in untreated WT bacteria; for RNA-seq, FDR using Benjamini-Hochberg.

### Denitrifying conditions in culture do not replicate *in planta* conditions

Because xylem fluid contains relatively small amounts of nutrients, minimal media has often been considered to mimic the xylem environment (Arlat et al. 1992; Lowe-Power et al. 2018a). Because VDM is a minimal medium and *R. solanacearum* experiences low-oxygen conditions and uses denitrifying respiration during tomato infection, we asked if anaerobic growth in VDM would better mimic the tomato xylem environment than rich media (Dalsing et al. 2015). However, when we examined expression of *R. solanacearum* genes in tomato stems compared to in VDM, we found a total of 1038 genes were differentially regulated in these conditions (Supplementary Table 2). Most genes involved in denitrification were expressed at a higher level in VDM than *in planta*. Other broad categories that were altered include sugar metabolism, flagellar motility, amino acid transporters, iron uptake genes, and ribosomal proteins. In line with previous results, T3SS-related genes were among the most highly upregulated genes *in planta* (Jacobs et al. 2012). Overall, growth in VDM did not closely replicate the *in planta* environment and the bacterium seemed more reliant on denitrifying metabolism in VDM.

### Altering denitrification and NO levels changes expression of denitrification and virulence-related genes *in planta*

To determine if the effects of NO on the T3SS were specific or a part of a non-specific general response, we examined genes differentially regulated the non-denitrifying *ΔnarG* and NO-accumulating *ΔnorB in planta*. In the stem, 187 genes were significantly altered by mutating *narG* and thus blocking denitrification and NO production, while only 49 genes were altered in the NO accumulating *ΔnorB* mutant. In *ΔnarG*, nine of the most significantly upregulated genes were directly involved in nitrate sensing and uptake or were components of the nitrate reductase protein. In both *ΔnarG* and *ΔnorB*, genes involved in motility and sulfur metabolism were decreased compared to WT (Supplementary Table 2). Overall, this muted response highlights the robust regulatory state of *R. solanacearum* cells growing *in planta* and shows that they can tolerate even changes in fundamental aspects of its biology (Peyraud et al. 2018).

Consistent with the observation that the entire transcriptome of WT *R. solanacearum* was very different *in planta* than in culture (Fig. 2aA) the transcriptomic profiles of the denitrification mutants grown in culture were much more divergent from that of WT, with 3,145 significantly altered genes in *ΔnarG* and 3,976 altered in *ΔnorB*. In the non-denitrifying *ΔnarG*, the bacterium showed decreased expression in many KEGG pathways involved in growth and metabolic activity such as purine and pyrimidine metabolism, oxidative phosphorylation, and tRNA synthetases (Supplementary Table 3).

In culture, genes up-regulated in the NO-accumulating *ΔnorB* mutant were enriched in the KEGG pathways of flagellar assembly, sulfur relay system and fatty acid degradation (Supplementary Table 3). Downregulated genes were enriched in central metabolic KEGG pathways like oxidative phosphorylation, pyrimidine metabolism, biosynthesis of amino acids, and fatty acid biosynthesis. In contrast to the activation of the T3SS, CysNO-treated *R. solanacearum* cells and *ΔnorB* saw reduction in expression of Types I, II, and VI secretion systems. Overall, the transcriptomic responses of *Rs* cells to either increased or decreased NO did not align well with their responses *in planta*. This is consistent with previous studies showing that cultured *Rs* cells do not reflect biology of the pathogen in its natural habitat (Jacobs et al. 2012; Perrier et al. 2018).

### *R. solanacearum* encounters more nitrosative stress in VDM than *in planta*

To estimate the degree of nitrosative stress induced by our treatments, we looked at the expression of *norB* and *hmpX*, which directly detoxify NO in *R. solanacearum* (Farr and Kogoma 1991; Dalsing et al. 2015). Compared with *R. solanacearum* growing in denitrifying culture conditions, *norB* and *hmpX* expression decreased 5-fold and 20-fold *in planta*, suggesting that in this environment the bacterium experiences lower nitrosative stress levels (Supplementary Table 2). In striking contrast, expression of the oxidative stress-response catalase gene *katE* was increased 35-fold *in planta. In planta*, neither the non-denitrifying *ΔnarG* strain nor the NO overproducing *ΔnorB* strain showed significant changes *norB* or *hmpX* expression. Conversely, the *ΔnorB* mutant grown in culture had an 11-fold up-regulation of the NO-detoxifying *hmpX*. In culture, our oxidative stress control treatment with H_2_O_2_ increased expression of *katE*, as expected, but also had increased expression of NO detoxifying genes *hmpX* and *norB*. Overall, these results showed some overlap in the response to ROS and RNS stress in culture but suggested that *R. solanacearum* cells experienced less nitrosative stress in tomato xylem than when denitrifying in VDM culture.

### Nitric oxide induced expression of T3SS-related genes both *in vitro* and *in planta*

To determine if in addition to *hrpB*, other T3SS-associated genes are induced by NO in culture, we measured relative expression of all 107 known *R. solanacearum* T3SS genes under each tested condition. These genes encode T3SS structural elements, regulators, secreted effectors, and chaperones (Fig. 4) (Cunnac et al. 2004; Peeters et al. 2013; Van Gijsegem et al. 1995; Plener et al. 2010; Lonjon et al. 2017; Zhang et al. 2015b, 2018, 2011; Brito et al. 1999; Marc Marenda et al. 1998; Lonjon et al. 2015; Génin et al. 2005). Cells of the NO-accumulating *ΔnorB* mutant grown in culture had broadly increased T3SS gene expression relative to wild type, with nine of sixteen structural elements significantly increased. Only one structural element, *hrpF*, trended non-significantly towards reduced expression (Fig. 4A). Five out of thirteen known regulators of T3SS were significantly upregulated in *ΔnorB*, although expression of the *prhA, prhG, prhO*, and the *prhKLM* operon were all reduced. Two out of the four T3E chaperones had significantly elevated expression in *ΔnorB*, as did most T3Es. The differential expression of effectors did not seem to correlate with their known interactions with chaperones (Supplementary Table 4). The expression of T3SS genes in CysNO-treated cells largely mirrored that of *norB*, with 80% of tested genes having a similar trend. However, the magnitude of the effect was lower. T3SS gene expression in *ΔnarG* was more varied, with fewer significantly differentially regulated genes than *ΔnorB*. In contrast to *ΔnorB, ΔnarG*, and CysNO-treated cultures, H_2_O_2_-treated cultures had only eight significantly changed T3SS genes (Fig. 4A). Overall, we saw a greater induction of T3SS genes in culture by our high-NO conditions than low-NO conditions.

**Fig 4.**
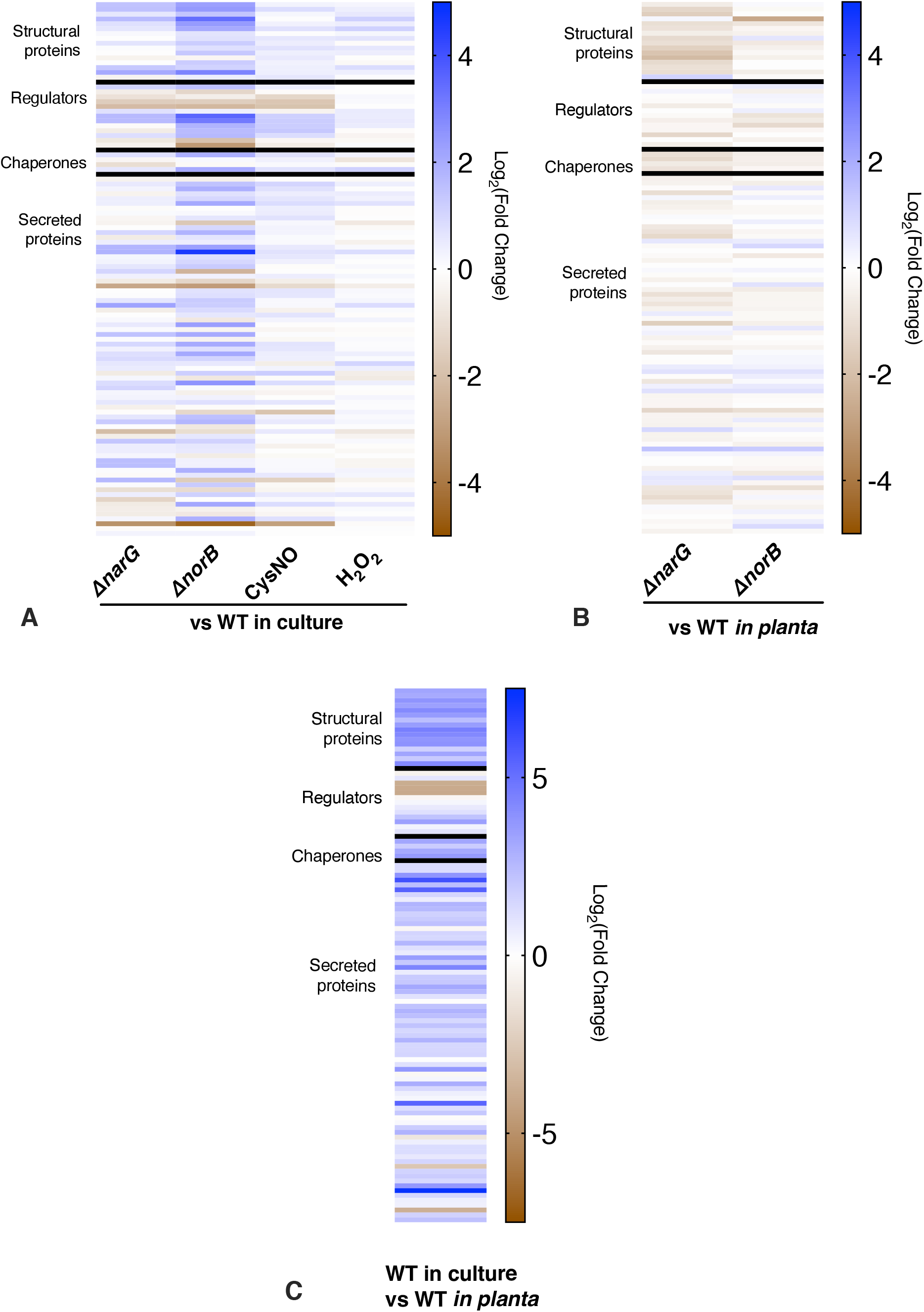
Nitric oxide alters expression of T3SS-related genes both *in vitro* and *in planta*. **A**, Heatmap showing relative expression levels of 107 T3SS-associated genes in *R. solanacearum* cells growing in rich medium with varying NO levels. All fold-change numbers are compared to levels in untreated wild-type (WT) cells. Gene annotations, fold change, and significance levels can be found in the supplementary materials. NO levels were modified genetically using mutants lacking *narG* (deleting this nitrate reductase reduces endogenous NO) and *norB* (deleting this nitric oxide reductase increases endogenous NO). NO levels were increased chemically by adding 1 mM CysNO. Cells treated with 100 µM hydrogen peroxide (H_2_O_2_) were a non-NO control to show general stress responses. **B**, Relative expression levels of 107 T3SS-associated genes in *R. solanacearum* cells growing *in planta*. About 2000 CFU of WT *R. solanacearum, ΔnarG*, or *ΔnorB* were introduced into 17-day old Bonny Best tomato plants through a cut petiole and RNA was harvested from a stem slice 3 days after inoculation. **C**, Expression of most T3SS-associated genes in *R. solanacearum* strain GMI1000 was enhanced when the bacterium grew in tomato plants relative to when *R. solanacearum* cells multiplied *in vitro* under denitrifying conditions (VDM medium, 0.1% O_2_).

To determine if altering NO levels affects *R. solanacearum* T3SS gene expression in a biologically relevant environment with all natural T3SS regulating inputs, we examined changes in T3SS gene expression in *ΔnarG* and *ΔnorB in planta*. Compared to *R. solanacearum* grown in culture, wild type *R. solanacearum in planta* had much higher expression of almost every T3SS-associated gene (Fig. 3C). When growing in tomato stems, *ΔnorB* grown *in planta* had no significantly differentially regulated T3SS genes, possibly because relatively high levels of NO are already present in the plant environment (Fig. 3B). In contrast, the non-denitrifying *ΔnarG* mutant showed an overall trend of lower T3SS gene expression, with expression of nine genes significantly reduced (Fig. 3B). These included two components of the T3SS structure, *hrpF* and *hrcN*, the regulators *hrpB* and *prhG*, the effector chaperone *hpaG*, and four T3SS effectors. Even in the highly T3SS-inducing *in planta* environment, reducing *R. solanacearum*-produced NO was sufficient to reduce T3SS gene expression in tomato xylem.

## Discussion

The T3SS is essential for *R. solanacearum* virulence, as it is for many Gram-negative pathogenic bacteria. Involving at least 16 structural proteins, this secretion system is metabolically expensive to synthesize and operate, so the bacterium is under strong selection pressure to express it only when it increases fitness (Genin et al. 1992; Boucher et al. 1985). Many factors contribute to the regulation of T3SS in *R. solanacearum*, including the presence of undefined host cell wall materials, unknown soluble factors, unknown metabolic cues, and (in culture) quorum sensing (Zuluaga et al. 2013; Brito et al. 1999; Aldon et al. 2000; Genin and Denny 2012; Zhang et al. 2018, 2011; Senuma et al. 2020). While many T3SS regulators have been described, open questions remain, such as why T3SS is repressed in rich media and whether specific host cell wall components activate PrhA. In addition, many genes that affect T3SS gene expression do so via unknown mechanisms (Zhang et al. 2011, 2015b, 2018). The results presented here suggest that NO participates in this complex regulatory network.

Despite its toxicity at high concentrations, NO is widely used as a signaling molecule in all domains of life (Asgher et al. 2017; Astuti et al. 2018; Simmonds et al. 2014). Many effects of NO are mediated by NO-dependent protein post-translational modifications, including binding transition metals, S-nitrosylation of cysteine, and tyrosine nitration (Simmonds et al. 2014; Cooper 1999; Leitner et al. 2009; Wünsche et al. 2011; Kolbert et al. 2017; Zhang et al. 2015a; Russwurm and Koesling 2004). While most well-characterized mechanisms of NO signaling have been described in eukaryotes, examples of NO signaling in bacteria have also been identified. Species in *Legionella, Shewanella, Vibrio*, and *Silicibacter* have been shown to use H-NOX proteins to sense NO and regulate biofilm formation (Rao et al. 2015; Henares et al. 2012; Liu et al. 2012; Network et al. 2012; Carlson et al. 2010). NO also contributes to biofilm regulation in *Pseudomonas aeruginosa* via the protein NosP (Hossain and Boon 2017). Both H-NOX proteins and NosP bind NO using iron centers, but no orthologues of these genes are apparent in the *R. solanacearum* genome. The first example of S-nitrosylation in bacteria was the oxidative stress response protein OxyR in *E. coli* (Seth et al. 2012). OxyR regulates a distinct set of genes when a key cysteine is nitrosylated rather than oxidized. In addition to this, nitrosative stress damage triggers S-nitrosylation of the *Salmonella enterica* redox sensor SsrB and affects the S-nitroso proteome of *Mycobacterium tuberculosis* (Rhee et al. 2005; Husain et al. 2010).

Although we show NO activates a key virulence factor in *R. solanacearum*, its specific mechanism of action is not yet known. Using *in silico* predictions of NO-dependent post-translational modifications, we could identify and selectively modify potentially important residues in T3SS regulatory proteins to determine the mechanism of NO-dependent T3SS activation (Xue et al. 2010; Liu et al. 2011).

We found that expression of T3SS-associated genes was enhanced by the presence of a plant host and by exogenous NO released by the NO donor CysNO, and the plant-mediated induction was suppressed by the NO scavenger cPTIO (Fig. 1C). T3SS gene expression also increased in response to nitric oxide produced by *R. solanacearum* itself, both at natural levels generated by denitrifying respiration and at enhanced levels caused by deleting *norB*. As previously reported (Monteiro et al. 2012a; Jacobs et al. 2012; Khokhani et al. 2017), *R. solanacearum* cells infecting tomato plants highly expressed most T3SS genes, but reducing *R. solanacearum*-produced NO by preventing denitrifying respiration with a Δ*narG* mutation significantly decreased expression of some T3SS-related genes *in planta*. These included genes encoding structural components, HrpF and HrcN, key regulators HrpB and PrhG, the chaperone HpaG, and four T3-secreted effectors (Fig. 4B). This was especially striking because overall gene expression in *ΔnarG* was otherwise very similar to that of wild type. These results raise the possibility that *R. solanacearum* uses the presence of NO as an indirect indicator of the host xylem environment. There was no increase in T3SS gene expression in the NO-overproducing *ΔnorB* mutant, which may indicate that T3SS genes are already so highly expressed *in planta* that the addition of more NO does not further increase this elevated level of expression.

Besides its effects on the T3SS, altering NO concentrations in culture elicited broad changes in gene expression. We reduced NO levels using the non-denitrifying mutant *ΔnarG*. However, preventing the cells from using the denitrification pathway has significant effects on the biology and metabolic state of the cells, making the interpretation of these results challenging. For example, the *ΔnarG* cells grown in culture showed evidence of decreases in growth rate and metabolic activity, indicated by decreased expression in the biosynthetic pathways of nucleotides, reduced expression of the oxidative phosphorylation system, and decreased production of many tRNAs. Treating the cells with NO, either with exogenously applied CysNO or with the NO-overproducing *ΔnorB*, caused similar changes in many metabolic pathways in culture. However, a primary force in culture was the stress caused by increased NO concentrations. This manifested in decreased expression of denitrification and nitrate uptake functions and an increased expression of the NO-detoxifying enzyme *hmpX*. This stress was associated with decreased expression of most *R. solanacearum* secretion systems. However, expression of the T3SS showed the opposite trend, indicating the specificity of NO-dependent T3SS regulation.

When denitrification mutants grew *in planta* they had far fewer alterations in gene expression relative to wild type than when the mutants grew in culture. This suggests that inputs from the plant host help *R. solanacearum* control its metabolism and behaviors. Overall, the cells grown *in planta* seem to encounter less nitrosative stress than cells grown in denitrifying conditions in culture. This could mean that NO stress is not as important *in planta* than in culture, but it does not take into account differences in the microenvironments of the xylem environment. Bacteria grown in culture experience a uniform environment with consistent NO stress. In contrast, *R. solanacearum* cells colonizing plant stems occupy multiple different microenvironments, which likely contain varying NO levels.

A previous study compared gene expression between *R. solanacearum* grown in rich media and *R. solanacearum* living in tomato xylem (Jacobs et al. 2012). Although there were methodological differences between that experiment and this one, we saw many similarities in the DEGs present between in culture and *in planta R. solanacearum*. In both experiments, many of the most highly upregulated genes are involved in the T3SS. Of the genes that were upregulated *in planta* in both experiments, 31 out of 141 (~22%) encoded structural components, regulators, or effectors of the T3SS. Three exceptions to this trend were *prhKLM*. Mutating any of these genes leads to a loss of T3SS gene expression, yet all three were highly downregulated *in planta* compared to either CPG or VDM (Supplementary Table 4) (Zhang et al. 2011; Jacobs et al. 2012). Several other genes that were highly upregulated *in planta* in both experiments encode metabolism of sucrose or myo-inositol, two carbon sources important for *R. solanacearum* during infection (Lowe-Power et al. 2018b; Hamilton et al. 2020; Xian et al. 2020). Additionally, genes involved in flagellar motility were upregulated in both experiments. A similar number of genes (137) were downregulated *in planta* in both experiments. These included several genes predicted to be involved in amino acid transport and metabolism and the superoxide dismutase genes *sodB* and *sodC*.

Components of the denitrification pathway were among the most highly upregulated *R. solanacearum* genes *in planta* compared to in rich medium. Consistent with this observation, *R. solanacearum* mutants unable to denitrify were reduced in growth *in planta* and in bacterial wilt virulence (Dalsing et al. 2015). However, the same genes were around 20-fold more highly expressed in denitrifying culture than *in planta*. Further, the RNS-responsive NO-detoxifying genes *norB* and *hmpX* were upregulated *in planta* compared to rich media but were expresses much less *in planta* than in denitrifying conditions in culture. This indicates that *Rs* experiences more nitrosative stress when growing in denitrifying culture conditions than when growing in the plant host, which in turn is more nitrosatively stressful than aerobic growth in rich media.

These data add to the strong evidence that *R. solanacearum* gene expression and metabolism are very different in the biologically realistic host environment than in culture. This may be due to the complex structure of the xylem environment, where host inputs modulate *R. solanacearum* gene expression, the ability to form defensive biofilms and multiple microenvironments with different nutrient or oxygen concentrations, or because the xylem is a flowing environment, constantly bringing new nutrients into the bacterial habitat. The bacterium’s behavior in culture should be regarded as artifactual unless proven otherwise.

We attempted to directly measure the concentrations of NO in healthy and infected tomato xylem collected from fluid pooling on a cut stem using the NO-reactive fluorescent dye DAF-FM, electro paramagnetic resonance (EPR), and an NO-sensitive microsensor (Namin et al. 2013; Kleschyov et al. 2007; Yoshioka et al. 1996; Schreiber et al. 2008). While a fluorescent signal was observed when DAF-FM was added to xylem sap, no signal was detected through EPR using the NO spin trap cPTIO or with the microsensor (Unisense, Aarhus, Denmark), suggesting that the DAF-FM fluorescence was not specific to NO. These results highlight the need to measure NO using multiple methods in biological systems (Gupta and Igamberdiev 2013). This failure to detect NO in tomato xylem may not reflect true levels of NO in an intact plant. We collected xylem sap by de-topping tomato plants just above the cotyledons and allowing root pressure to force xylem fluid to pool on the cut stem. Any material exuded in the first two minutes after cutting was removed in an attempt to limit the amount of cellular debris and phloem in the collected material. This method, while useful for obtaining sap to analyze for stable metabolites may not work with a transient metabolite like NO (Lowe-Power et al. 2018a). Cutting the stems stimulates production of reactive oxygen species, which quickly react with NO to form other reactive nitrogen species, potentially diluting any NO signal (Milling et al. 2011; Beckman and Koppenol 1996; Van Faassen and Vanin 2007). Furthermore, this approach can only measure NO levels in bulk xylem sap. The xylem is a dynamic and physically heterogenous environment. Xylem flow, xylem structure, and *R. solanacearum*-produced biofilms may all create gradients or microenvironments with altered NO concentrations in the xylem of intact plants (Zimmermann, 1983; Kim *et al*., 2016; Tran *et al*., 2016; Lowe-Power, Khokhani and Allen, 2018). These factors complicate direct measurement of xylem NO. Confocal microscopy of intact infected stems using fluorescent markers to visualize *R. solanacearum*, biofilms, and NO may eventually give a more complete understanding of this challenging landscape.

Overall, these results reveal a strong connection between NO levels and expression of the *R. solanacearum* T3SS. They suggest that *R. solanacearum* regulates virulence in part by using denitrifying metabolism and the resulting NO as a signal of the hypoxic high-nitrogen host xylem environment. Ongoing work is exploring how *R. solanacearum* manages nitrosative stress and identify additional effects of NO on biology of both host plant and bacterial pathogen.

## Materials and Methods

### Bacterial strains and culture conditions

All strains, plasmids, and primers used in this study are listed in Supplementary Table 1. *R. solanacearum* was cultured in either CPG or a modified VDM medium at 28°C (Hendrick and Sequeira 1984; Dalsing et al. 2015). *E. coli* was grown in Luria-Bertani (LB) media at 37°C. As needed, cultures were supplemented with antibiotics: 25 μg/mL kanamycin or 15 μg/mL gentamicin.

### Strain construction

An *R. solanacearum* GMI1000 mutant lacking *hrpB* (RSp0873) was created using a modified *sacB*-dependent positive selection vector, pUFR80 (Castañeda et al. 2005). The regions directly upstream and downstream of *hrpB* were amplified using the ΔhrpBup/dwn primer pairs. These fragments were then inserted into pUFR80 digested with *Hind*III using Gibson assembly (Sievers et al. 2011). The resulting plasmid, pUFR80-ΔhrpB was transformed into *R. solanacearum* GMI1000 and successful plasmid integrations were selected for kanamycin resistance. The resulting colonies were then counter selected on CPG + 5% w/v sucrose. Successful deletions were confirmed using PCR, sequencing, and a functional screen for the ability to induce HR on the incompatible host *N. tabacum* (Poueymiro et al. 2009). The *ΔhrpB* mutation was complemented by inserting a DNA fragment encoding *hrpB* and *hrcC* with the ~600 bp upstream containing their native promoter into the selectively neutral *att* site in the *R. solanacearum* chromosome using pRCK (Monteiro et al. 2012b). The fragment was amplified using hrpBcomp_F/R and inserted into *AvRII/XbaI*-digested pRCK using Gibson assembly. This vector was then transformed into *ΔhrpB* and selected on CPG+kan.

### Plant growth conditions

Wilt-susceptible tomato cv. Bonny Best were germinated and grown in BM2 all-purpose germination and propagation mix (Berger, Saint-Modeste, QC). Plants were grown at 28°C with a 12 hr photoperiod. Seedlings were transplanted after 14 days. After transplanting, plants were watered on alternate days with 1/2 strength Hoagland solution. Tomato seeds for axenic seedling production were sterilized by washing with 10% bleach for ten minutes, followed by two five-minute washes in 70% ethanol, then rinsed five times with water and germinated on water agar plates.

### Lux reporter assays

To test the effect of denitrification on T3SS gene expression, *R. solanacearum* strain GMI1000 carrying a *hrpB::lux* reporter gene fusion was grown in denitrifying conditions as follows (Monteiro et al. 2012a). Overnight cultures grown in CPG were resuspended in VDM with or without 30 mM potassium nitrate to an initial OD_600_ of These cultures were grown in 200 µL volumes in clear-bottomed white-walled 96 well plates at 28°C in 0.1% O_2_ with shaking. Every half hour, the Abs_600_ and luminescence was measured using a BioTek SynergyHT microplate reader (BioTek, Winooski, VT, USA). To determine the effects of the NO scavenger cPTIO on T3SS gene expression, *hrpB::lux* cultures were resuspended in 24-well plates in 1 mL of water, either with or without two five day-old sterile tomato seedlings. As appropriate, cPTIO was added to a final concentration of 1 mM. The plates were incubated at 28°C with shaking for ~24 hours. The pigmented liquid was removed from the seedlings and transferred to 1.5 mL Eppendorf tubes, after which the bacteria were pelleted by centrifugation. The pigmented supernatant was removed, and the pellets resuspended in 200 µL of water. These suspensions were transferred to a new plate and the Abs_600_ and luminescence was measured as detailed above.

### RNA extraction for sequencing

To profile gene expression of *R. solanacearum* at various levels of NO stress, we extracted RNA from *R. solanacearum* grown in culture and *in planta*. For each in condition, we used *R. solanacearum* from three separate overnight cultures as biological replicates. For the *in vitro* samples, overnight cultures grown in CPG were resuspended to a final OD_600_ of 0.01 in 50 mL of VDM + 30 mM potassium nitrate in conical tubes. These cultures were incubated without shaking for 12 hours at 28°C in 0.1% O_2_. Four hours before harvesting, CysNO or H_2_O_2_ were added to the relevant cultures to a final concentration of 1 mM and 100 μM, respectively. CysNO was synthesized by adding HCl to a solution containing 200 mM L-cysteine and 200 mM sodium nitrite. The solution was incubated in the dark for ten minutes before it was neutralized by adding sodium hydroxide to a final concentration of 200 mM. The molarity of the solution was determined using the absorbance at 334 nm and the extinction coefficient 90/M*cm. After 16h total incubation, sub-samples were collected for dilution plating to determine CFU/ml and the tubes were capped and centrifuged at room temperature for 5 min at 8000 rpm. The supernatant was removed and pellets were frozen in liquid nitrogen. RNA extractions were carried out using a modified version of the Quick-RNA™ MiniPrep kit (Zymo Research, Irvine, CA, USA), as follows. Pellets were resuspended in 400 µL of ice cold TE pH 8 with 1 mg/mL lysozyme, 0.25 µL Superase Inhibitor (Ambion, Austin, TX, USA), and 80 µL of 10% SDS, vortexed for 10s, transferred to a new 2 mL tube, and shaken at ~300 rpm for 2 min. 800 µL of RNA-Lysis lysis buffer was added, the tubes were vortexed again for 10 s, cleaned according to the kit manufacturer’s instructions, and eluted in 100 µL of water. Nucleic acid concentrations were estimated using a nanodrop, normalized to 200 ng/uL, and cleaned using the DNA-free DNAse kit (Invitrogen, Carlsbad, CA, USA). Samples were incubated at 37C for 1 hour, with 2 µL more of DNAse added at 30 min. After DNAse inactivation, samples were further cleaned by chloroform extraction, then precipitated overnight at −20°C with 100 *μ*M Sodium Acetate pH 5.5 and 66 % ethanol. Samples were checked for concentration on a Nanodrop, for DNA contamination by PCR using the qRT-PCR primers *serC*_F/R, and for RNA integrity (RIN) using an Agilent Bioanalyzer 21000 (Agilent, Santa Clara, CA, USA). All samples had RIN values above 7.3.

The *in planta* samples were harvested after 21 day old Bonny Best tomatoes were inoculated with ~2000 CFU of each bacteria strain through the cut petiole of the first true leaf. At 3 dpi, approximately 0.1 g of stem tissue was collected from the site of inoculation, immediately frozen in liquid nitrogen, and stored at −80°C. Another ~0.1 g of tissue was collected from directly below the inoculation site and was ground in bead beater tubes using a PowerLyzer (Qiagen, Hilden, Germany) for two cycles of 2200 rpm for 90 s in each with a 4 min rest between cycles. This material was then dilution plated to measure bacterial colonization. From each biological replicate, we chose six plants with the most similar level of colonization. RNA was extracted from these using a modified hot phenol-chloroform method (Jacobs et al. 2012). Between 4 and 5 individual plants were pooled per biological replicate. Nucleic acid sample quality was checked using a nanodrop, Agilent bioanalyzer, and qRT-PCR primers Actin_F/R. All samples had RIN values of above 7.2.

### RNA sequencing and data analysis

All RNA was sent to Novogene (Beijing, China) for library preparation, sequencing, and analysis. rRNA depletion, fragmentation, and library construction were done using NEBNext® µLtra™ Directional RNA Library Prep Kit for Illumina® (NEB, Ipswitch, MA, USA) following the manufacturer’s recommendation and starting with 3 μg of RNA per sample. After the libraries were constructed, their quality was assessed using an Agilent Bioanalyzer 21000 system and sequenced using an Illumina platform to generate paired-end reads. Between 19,000,000 and 40,000,000 reads were produced per sample. Raw reads were converted to FASTQ files using Illumina CASAVA v1.8, and then filtered for quality, removing reads with adaptor contamination, greater than 10% uncertain nucleotides, or more than 50% of nucleotides with a Q_pred_ less than or equal to 5. Over 95% of reads were of good quality, and were mapped to the *R. solanacearum* GMI1000 genome (https://www.ncbi.nlm.nih.gov/assembly/GCF_000009125.1) using Bowtie2 −2.2.3 (Langmead et al. 2009). From this, gene expression was calculated using HTseq v0.6.1 and differential gene expression was calculated using DESeq 1.18.0. P-values calculated by DESeq were adjusted to control for the false discovery rate (FDR) using the Benjamini-Hochberg approach. Gene Ontology enrichment was analyzed using GOseq Release2.12 and KEGG enrichment was calculated using KOBAS v2.0 software.

### Measuring T3SS expression with qRT-PCR

Expression of T3SS genes was measured in denitrification mutants grown in 40 mL of VDM + 30 mM nitrate grown in GasPak™ EZ microaerobic pouches (BD, Franklin Lakes, NJ, USA) at 28°C with shaking for 18 hours starting from an OD_600_ of 0.001. RNA extractions were carried out using a hot phenol-chloroform method (Jacobs et al. 2012). Extractions were carried out using the Quick-RNA™ MiniPrep kit (Zymo Research, Irvine, CA, USA). cDNA synthesis was done using the Superscript VILO cDNA synthesis kit (Invitrogen, Carlsbad, CA, USA). qPCR reactions were carried out in 10 µL volumes using 5 ng of total template using PowerUp Syber Green Master) in a QuantStudio 5 Real-Time PCR System Mix (Applied Biosystems, Foster City, CA, USA).

## Data Availability

The data supporting this research are openly available in the Gene Expression Omnibus at https://www.ncbi.nlm.nih.gov/geo/query/acc.cgi?acc=GSE160024, GEO accession: GSE160024.

## Acknowledgments

The authors would like to thank Jon Jacobs for discussion and advice throughout the project and Max Miao for his expertise and discussion about statistics and R. CGH and BLD were supported by NSF predoctoral fellowships. This material is based upon work supported by the National Science Foundation Graduate Research Fellowship Program under Grant No. DGE-1747503. Any opinions, findings, and conclusions or recommendations expressed in this material are those of the author(s) and do not necessarily reflect the views of the National Science Foundation. The authors declare no competing financial interests

---

**Supplementary Table 2** Complete list of gene expression and log_2_ fold change values for all conditions included in this study. All fold-change numbers are compared to levels in untreated wild-type (WT) cells. NO levels were modified genetically using mutants lacking *narG* (deleting this nitrate reductase reduces endogenous NO) and *norB* (deleting this nitric oxide reductase increases endogenous NO). NO levels were increased chemically by adding 1 mM CysNO. Cells treated with 100 µM hydrogen peroxide (H_2_O_2_) were a non-NO control to identify general stress responses. *P*-values were calculated by DESeq were adjusted to control for the false discovery rate (FDR) using the Benjamini-Hochberg approach.

**Supplementary Table 3** KEGG enrichment of genes altered in each condition. enrichment was calculated using KOBAS v2.0 software. *P* values were calculated using a hypergeometric test and adjusted using the Benjamini and Hochberg FDR correction method.

**Supplementary Table 4** Annotations, Log_2_ fold change relative to wild-type and adjusted p-values for *R. solanacearum* T3SS related genes in response to changes in NO concentration both *in planta* and in culture (see Figure 3 for heatmap of these data). All fold-change numbers are compared to levels in untreated wild-type (WT) cells. *in planta* samples are denoted by a ‘p’ in front of the treatment (e.g. *pnorB*). Genes organized by their category, including structural genes, regulatory genes, chaperones, and effectors. Effectors are subgrouped based on their known association with chaperones (Lonjon et al. 2015).

**Supplementary Table 1.**
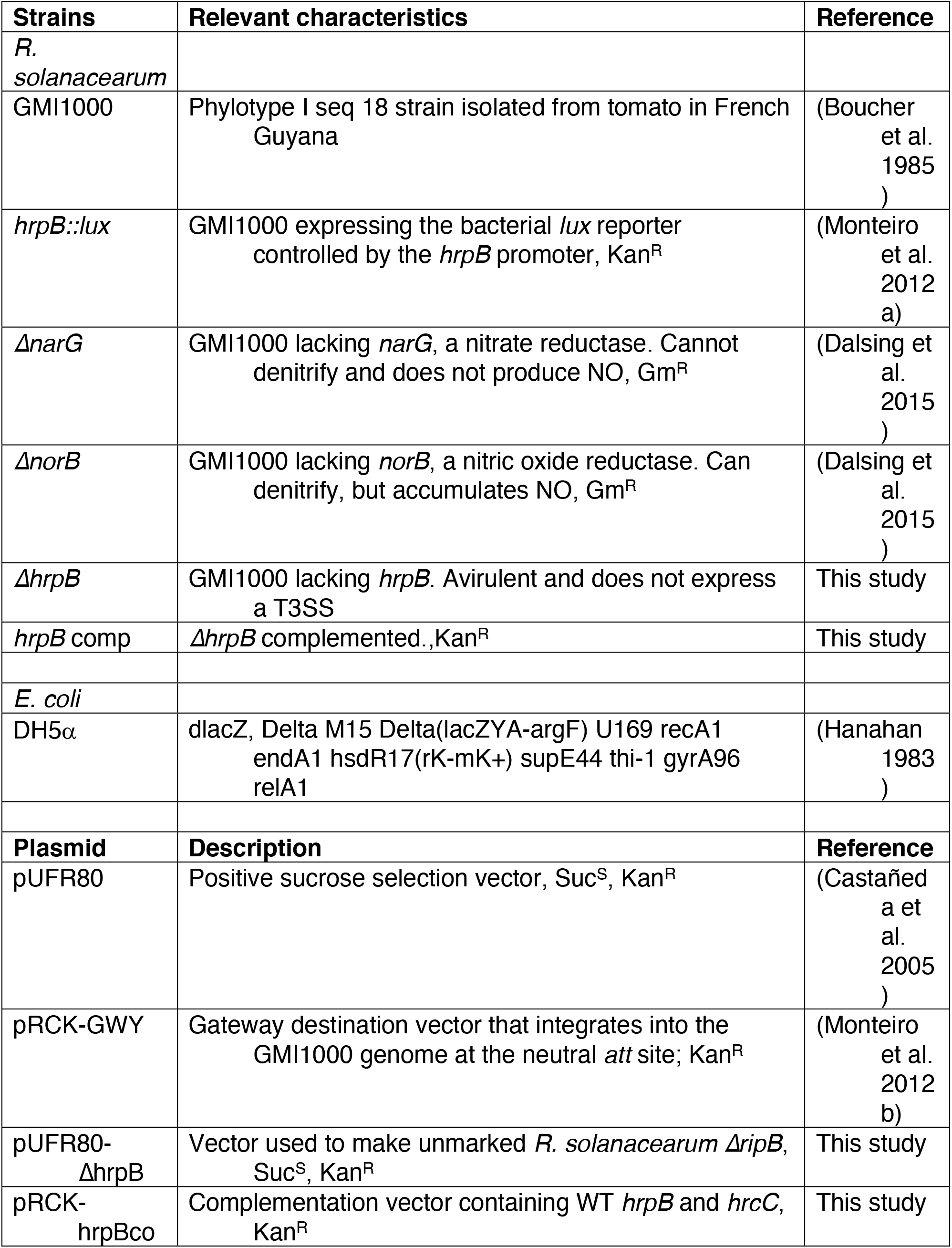
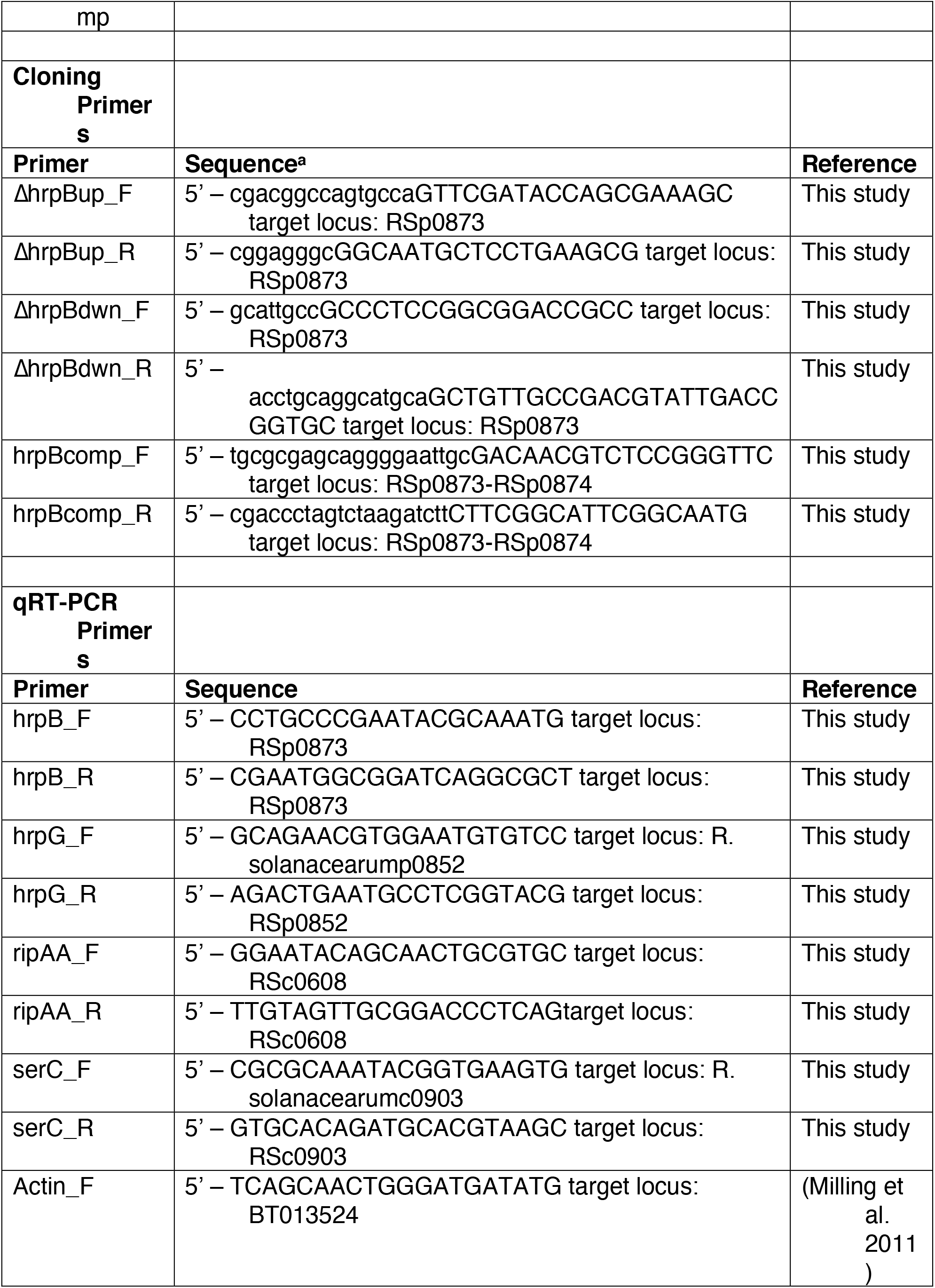
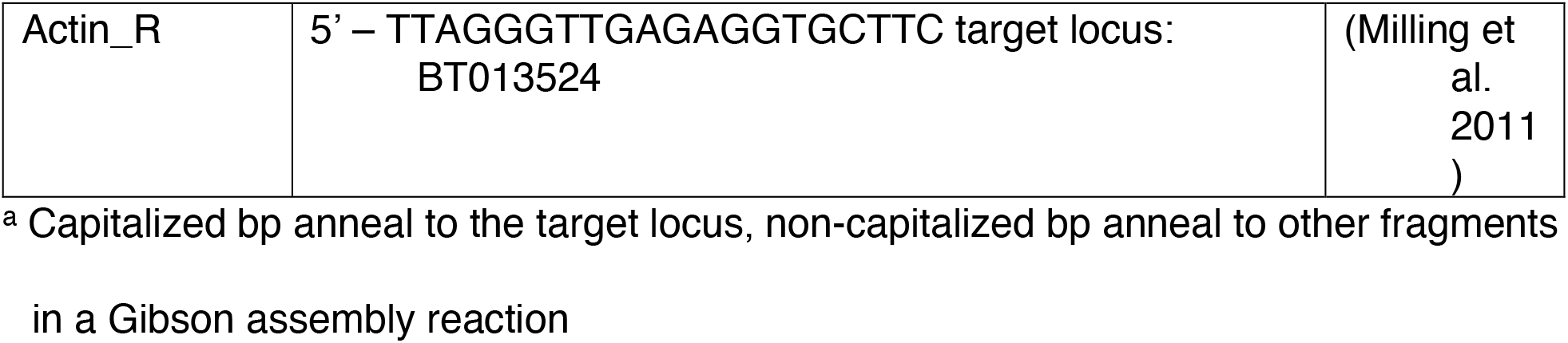
Bacterial strains, plasmids, and primers.

**Supplementary Fig 1.**
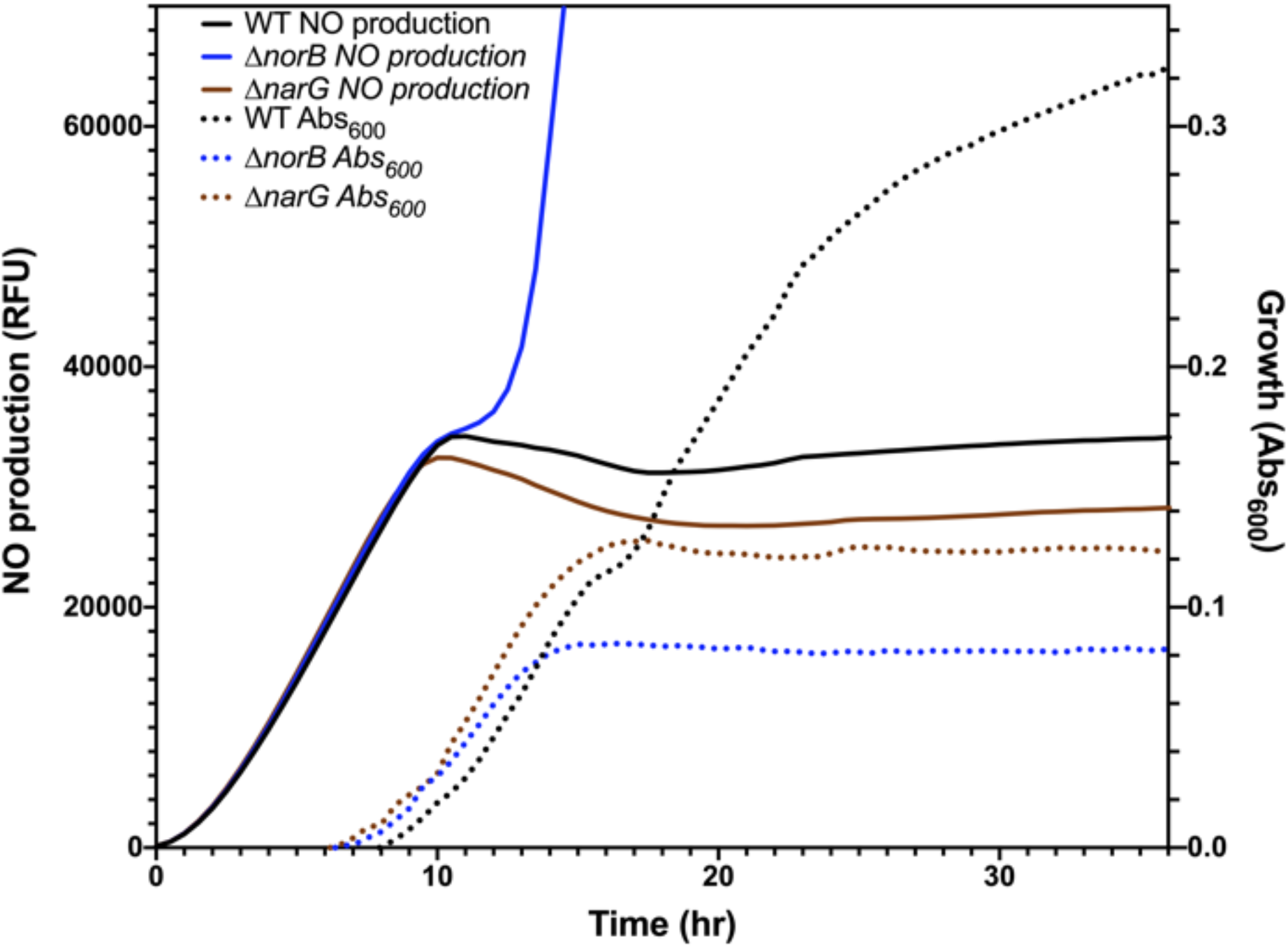
Growth and NO levels of *R. solanacearum* strains under denitrifying conditions in culture. Cultures were started at a concentration of 10^6^ CFU/mL in VDM containing the NO-detecting fluorescent dye DAF-FM DA at 10 µM. Cultures were incubated in in clear-bottomed, white-walled 96-well plates at 30C for 24 h at [O_2_] =0.1%. Culture growth (dotted lines) is shown as absorbance at 600 nm. NO concentration (solid lines) was measured as DAF-FM fluorescence at 495/515 nm. The *ΔnorB* mutant strain accumulates NO, which significantly impairs its growth.

**Supplementary Fig 2.**
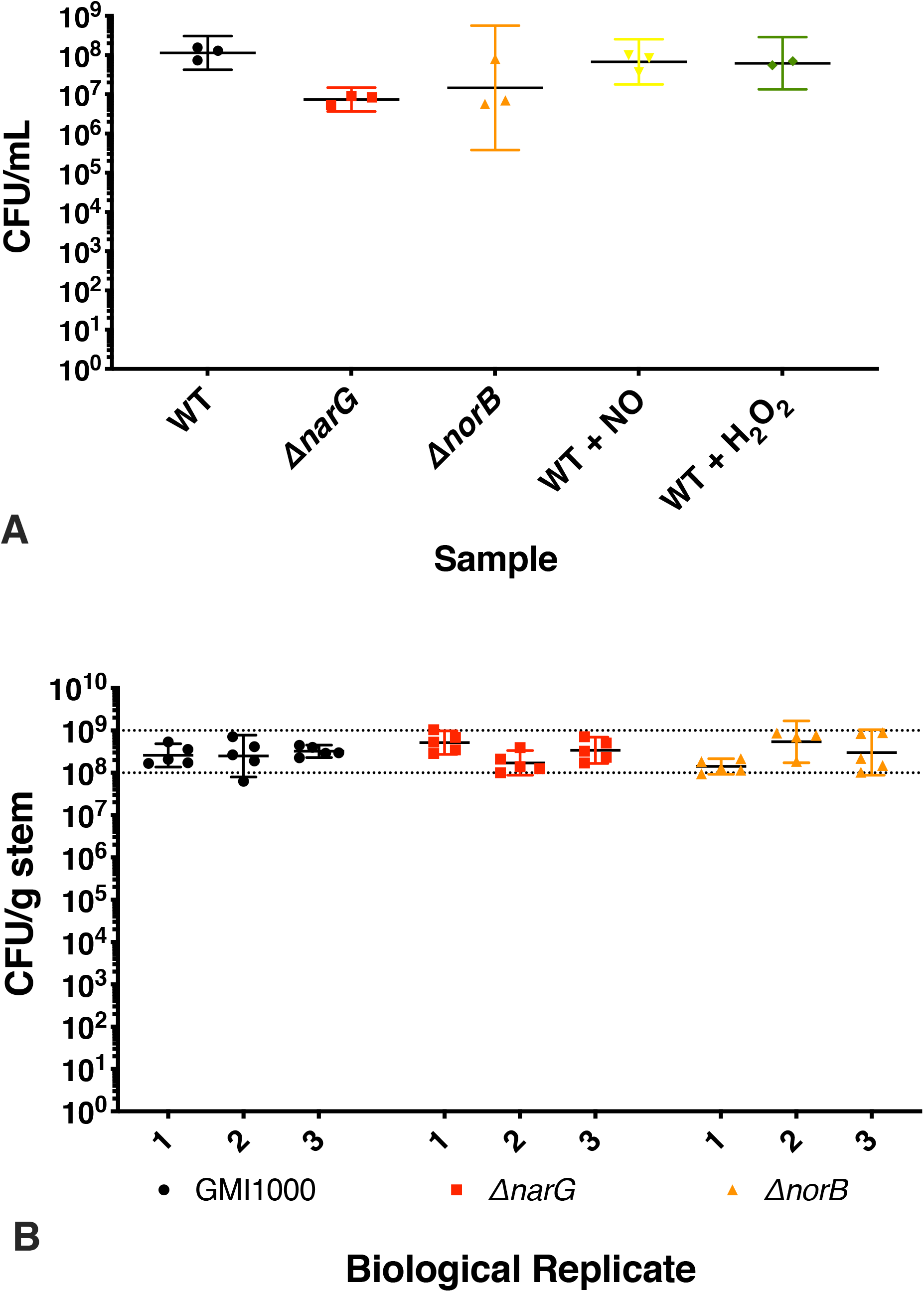
In vitro cell densities and tomato stem colonization of samples used in RNA-seq. **A**, Cell densities of culture samples as measured by dilution plating. One H_2_O_2_-treated plate was uncountable due to plate contamination. OD_600_ values of each culture were also collected and had comparable values. **B**, Colonization of tomato plants used for transcriptome sequencing. Roughly 0.1 g of stem tissue was collected from directly below the sample used for sequencing. The tissue was ground and dilution plated to enumerate CFU/g stem.

**Supplementary Fig 3.**
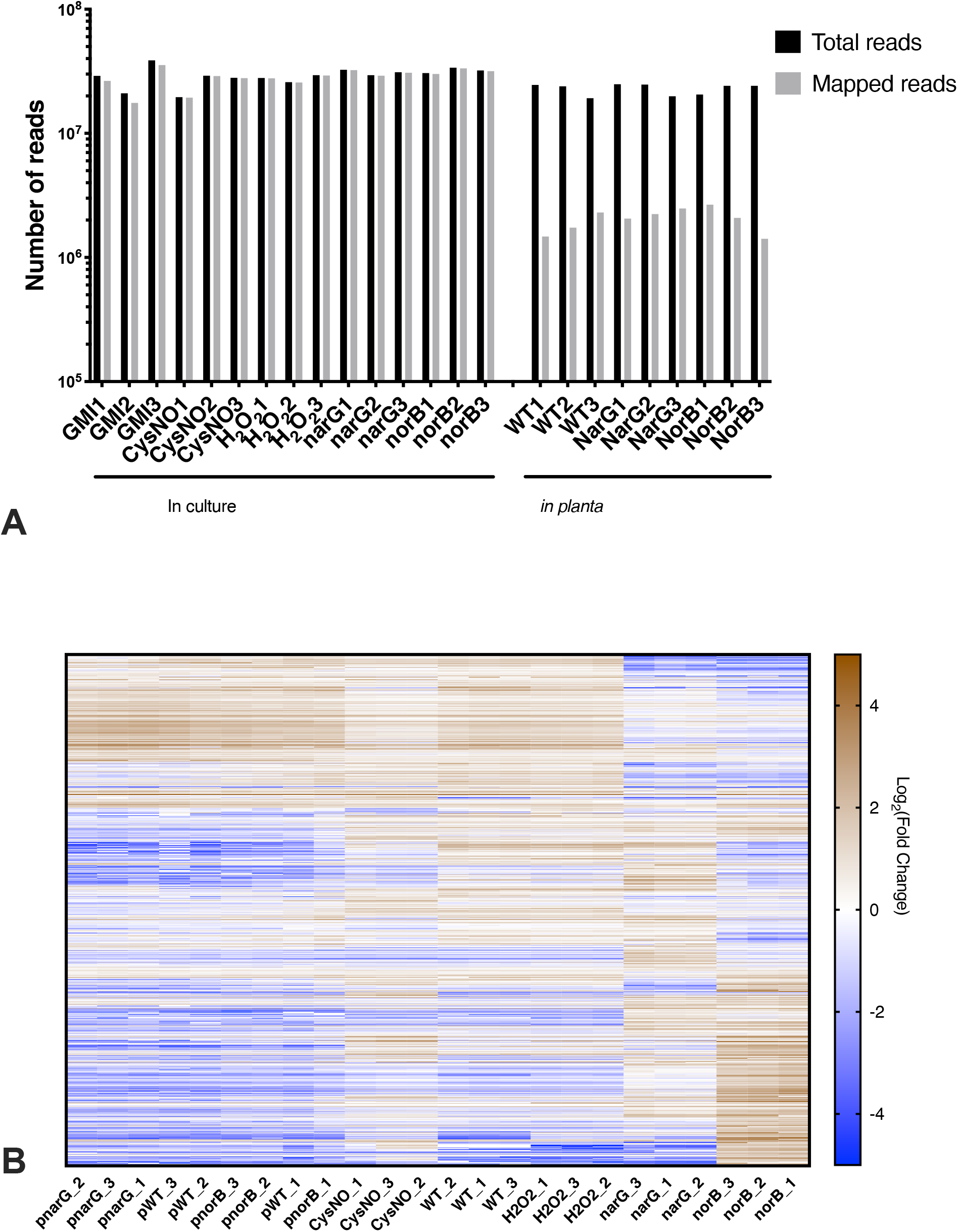
General RNA-seq quality **A**, Total and mapped reads per RNA-seq sample. The *in planta* samples had lower percentage of mapped reads due to the presence of host RNA. **B**, Hierarchical clustering and expression heat map of the 1000 most variable genes in the dataset. Blue indicates higher expression and brown indicates lower expression. All genes were normalized to the mean gene expression across all conditions. Samples collected from *R. solanacearum* grown *in planta* are denoted with the prefix “p”. Heatmap created using iDEP.90 (Ge et al. 2018).

